# Immunotherapy prevents long-term disability in relapsing multiple sclerosis over 15 years

**DOI:** 10.1101/735662

**Authors:** Tomas Kalincik, Sifat Sharmin, Charles Malpas, Tim Spelman, Dana Horakova, Eva Kubala Havrdova, Maria Trojano, Guillermo Izquierdo, Alessandra Lugaresi, Alexandre Prat, Marc Girard, Pierre Duquette, Pierre Grammond, Vilija Jokubaitis, Anneke van der Walt, Francois Grand’Maison, Patrizia Sola, Diana Ferraro, Vahid Shaygannejad, Raed Alroughani, Raymond Hupperts, Murat Terzi, Cavit Boz, Jeannette Lechner-Scott, Eugenio Pucci, Vincent Van Pesch, Franco Granella, Roberto Bergamaschi, Daniele Spitaleri, Mark Slee, Steve Vucic, Radek Ampapa, Pamela McCombe, Cristina Ramo-Tello, Julie Prevost, Javier Olascoaga, Edgardo Cristiano, Michael Barnett, Maria Laura Saladino, Jose Luis Sanchez-Menoyo, Suzanne Hodgkinson, Csilla Rozsa, Stella Hughes, Fraser Moore, Cameron Shaw, Ernest Butler, Olga Skibina, Orla Gray, Allan Kermode, Tunde Csepany, Bhim Singhal, Neil Shuey, Imre Piroska, Bruce Taylor, Magdolna Simo, Carmen-Adella Sirbu, Attila Sas, Helmut Butzkueven, on behalf of the MSBase Study group

**Affiliations:** CORe, Department of Medicine, University of Melbourne, Melbourne, Australia; Department of Neurology, Royal Melbourne Hospital, Melbourne, Australia; CORe, Department of Medicine, University of Melbourne, Melbourne, Australia; Department of Medicine, University of Melbourne, Melbourne, Australia; Department of Neurology, Royal Melbourne Hospital, Melbourne, Australia; Department of Neurology and Center of Clinical Neuroscience, General University Hospital and Charles University in Prague, Prague, Czech Republic; Department of Basic Medical Sciences, Neuroscience and Sense Organs, University of Bari, Bari, Italy; Hospital Universitario Virgen Macarena, Sevilla, Spain; Department of Neuroscience, Imaging and Clinical Sciences, University “G. d’Annunzio”, Chieti, Italy; Department of Biomedical and Neuromotor Sciences, University of Bologna; IRCCS Istituto delle Scienze Neurologiche di Bologna; Hopital Notre Dame, Montreal, Canada; CHUM and Universite de Montreal, Montreal, Canada; CISSS Chaudière-Appalache, Levis, Canada; Central Clinical School, Monash University, Melbourne, Australia; Neuro Rive-Sud, Quebec, Canada; Department of Neuroscience, Azienda Ospedaliera Universitaria, Modena, Italy; Isfahan University of Medical Sciences, Isfahan, Iran; Amiri Hospital, Kuwait City, Kuwait; Zuyderland Ziekenhuis, Sittard, Netherlands; Medical Faculty, 19 Mayis University, Samsun, Turkey; KTU Medical Faculty Farabi Hospital, Karadeniz Technical University, Trabzon, Turkey; School of Medicine and Public Health, University Newcastle, Newcastle, Australia; Department of Neurology, John Hunter Hospital, Newcastle, Australia; UOC Neurologia, Azienda Sanitaria Unica Regionale Marche - AV3, Macerata, Italy; Cliniques Universitaires Saint-Luc, Brussels, Belgium; University of Parma, Parma, Italy; C. Mondino National Neurological Institute, Pavia, Italy; Azienda Ospedaliera di Rilievo Nazionale San Giuseppe Moscati Avellino, Avellino, Italy; Flinders University, Adelaide, Australia; Westmead Hospital, Sydney, Australia; Nemocnice Jihlava, Jihlava, Czech Republic; University of Queensland, Brisbane, Australia; Royal Brisbane and Women’s Hospital, Brisbane, Australia; Hospital Germans Trias i Pujol, Badalona, Spain; CSSS Saint-Jérôme, Saint-Jerome, Canada; Hospital Universitario Donostia, Paseo de Begiristain, San Sebastián, Spain; Hospital Italiano, Buenos Aires, Argentina; Brain and Mind Centre, University of Sydney, Sydney, Australia; INEBA - Institute of Neuroscience Buenos Aires, Buenos Aires, Argentina; Hospital de Galdakao-Usansolo, Galdakao, Spain; Liverpool Hospital, Sydney, Australia; Jahn Ferenc Teaching Hospital, Budapest, Hungary; Craigavon Area Hospital, Craigavon, United Kingdom; Jewish General Hospital, Montreal, Canada; Deakin University, Geelong, Australia; Monash Medical Centre, Melbourne, Australia; South East Trust, Belfast, United Kingdom; Perron Institute, University of Western Australia, Nedlands, Australia; Institute of Immunology and Infectious Diseases, Murdoch University, Perth, Australia; Sir Charles Gairdner Hospital, Perth, Australia; Department of Neurology, Faculty of Medicine, University of Debrecen, Debrecen, Hungary; Bombay Hospital Institute of Medical Sciences, Mumbai, India; St Vincents Hospital, Fitzroy, Melbourne, Australia; Veszprém Megyei Csolnoky Ferenc Kórház zrt., Veszprem, Hungary; Royal Hobart Hospital, Hobart, Australia; Semmelweis University Budapest, Budapest, Hungary; Central Military Emergency University Hospital, Bucharest, Romania; BAZ County Hospital, Miskolc, Hungary; Department of Medicine, University of Melbourne, Melbourne, Australia; Central Clinical School, Monash University, Melbourne, Australia

**Author notes:** The list of MSBase contributors is provided in online Supplementary Table 1. *Statistical analysis* was completed by Tomas Kalincik of the University of Melbourne. **Corresponding author:** Tomas Kalincik, CORe, Level 4 Centre Royal Melbourne Hospital, 300 Grattan St, Melbourne VIC 3050.

**Keywords:** [41] Multiple sclerosis, [23] Clinical trials Observational study (Cohort, Case control), [54] Cohort studies, [324] Class III

## Abstract

**Objective:** Whether immunotherapy improves long-term disability in multiple sclerosis has not been satisfactorily demonstrated. This study examined the effect of immunotherapy on long-term disability outcomes in relapsing-remitting multiple sclerosis.

**Methods:** We studied patients from MSBase followed for ≥1 year, with ≥3 visits, ≥1 visit per year and exposed to a multiple sclerosis therapy, and a subset of patients with ≥15-year follow-up. Marginal structural models were used to compare the hazard of 12-month confirmed increase and decrease in disability, EDSS step 6 and the incidence of relapses between treated and untreated periods. Marginal structural models were continuously re-adjusted for patient age, sex, pregnancy, date, disease course, time from first symptom, prior relapse history, disability and MRI activity.

**Results:** 14,717 patients were studied. During the treated periods, patients were less likely to experience relapses (hazard ratio 0.60, 95% confidence interval 0.43–0.82, p=0.0016), worsening of disability (0.56, 0.38-0.82, p=0.0026) and progress to EDSS step 6 (0.33, 0.19-0.59, p=0.00019). Among 1085 patients with ≥15-year follow-up, the treated patients were less likely to experience relapses (0.59, 0.50–0.70, p=10^-9^) and worsening of disability (0.81, 0.67-0.99, p=0.043).

**Conclusions:** Continued treatment with multiple sclerosis immunotherapies reduces disability accrual (by 19-44%), the risk of need of a walking aid by 67% and the frequency of relapses (by 40-41%) over 15 years. A proof of long-term effect of immunomodulation on disability outcomes is the key to establishing its disease modifying properties.

## Introduction

Prevention of long-term disability accrual is currently the main goal of multiple sclerosis (MS) treatment. The available immunotherapies mitigate clinical and subclinical inflammation within the central nervous system.^1^ Some of these therapies reduce disability accrual over the short-term (≤3 years).^2–5^ Extension studies and randomised clinical trials suggested that timely immunotherapy may delay conversion to clinically definite MS,^6–8^ accumulation of disability^9, 10^ and death.^11^ However, observational studies reported conflicting results. One study did not find differences in disability outcomes between interferon β and no treatment (even though contrasting trends were seen when interferon β was compared to historical and contemporary untreated controls).^12^ Conversely, interferon β and glatiramer acetate were shown to mitigate disability accrual over ten years in the UK MS Risk Sharing scheme.^13, 14^

A proof of long-term effect of immunomodulation on the accumulation of MS-related neurological disability is the key to establishing its disease modifying properties. However, conclusive evidence is still lacking and it is unlikely that it will arise from randomised trials.

Here we present results from the largest international observational MS cohort, whose aim was to compare worsening and improvement of disability and incidence of relapses during periods of treatment vs. no treatment with MS immunotherapies over more than 15 years of follow-up. We hypothesised that continued treatment is associated with substantially reduced hazard of disability worsening over the long term. Comparisons of observational data between treated and untreated patients are obfuscated by strong indication bias, and randomised clinical trials that would address this question are neither feasible nor ethical. Therefore, this study used observational data analysed with marginal structural models to adjust for time-dependent confounding of treatment allocation.

## Methods

### Study design

This study compared long-term disability outcomes during periods under treatment and periods not under treatment recorded in an observational cohort of patients with MS and eligible for immunotherapy. The study estimated frequencies of relapses and disability accumulation or improvement events in patients who were hypothetically always exposed vs. never exposed to disease modifying therapies for MS. Because these two extreme scenarios are rarely directly observed, and if so, outcomes are usually strongly confounded by indication bias, we have used counterfactual framework to estimate causal associations between long-term exposure to therapies and outcomes (confirmed worsening or improvement of disability, incidence of relapses, EDSS step 6) based on the observed periods under treatment and not under treatment recorded in a single cohort. The counterfactual framework enables an analyst to quantify the probability of reaching disease outcomes under hypothetical conditions when the observed cohorts would remain always treated vs. never treated for the full duration of the follow-up period (“pseudo-cohorts”; Supplementary Figure 1 and 5), and continuously re-adjusted for confounders of outcomes with a marginal structural model.^15^

### Standard approvals

The MSBase registry is an international observational MS cohort, with contribution mainly from academic MS centres, registered with the WHO International Clinical Trials Registry Platform [ACTRN12605000455662].^16, 17^ The study was approved by the Melbourne Health Human Research Ethics Committee and the site institutional review boards. Patients provided written informed consent, as required.

### Patients

The inclusion criteria for this study were clinically isolated syndrome or definite MS.^18, 19^ The minimum required data consisted of follow-up ≥1 year, ≥3 disability scores with ≥1 score recorded per year, a minimum dataset (to enable evaluation of the outcomes and adjustment for confounders (see “Procedures”)) and exposure to a MS immunotherapy during the recorded follow-up. This was to exclude patients with benign disease course, who are unlikely to be treated, and to assure that only contemporary controls are included.

### Procedures

The data were recorded prospectively as part of clinical practice mainly at academic MS centres, as governed by the MSBase Observational Plan. Rigorous automated quality assurance procedure was applied (Supplementary Table 3).^17^

The follow-up time was segmented into 3-month periods (to maximise the use of the information from patients followed more frequently than the median visit frequency), with potential confounders and intermediates of treatment effect captured in MSBase at each period (for a causal diagram see Supplementary Figure 2). These consisted of time-dependent variables: treatment status (treated/untreated), treatment status during the preceding period, pregnancy status, pregnancy status during the preceding period, date of the period end, patient age, disease duration from the first MS symptom, disability score, change in disability score during the preceding 3 and 12 months, number of relapses during the preceding 3 and 12 months, the numbers of severe relapses, relapses with poor recovery and on-treatment relapses during the preceding 12 months, and MRI activity during the preceding 12 months; and fixed variables: sex, date of birth, date of first MS symptom, disease duration at first visit. Only periods with MS disease modifying therapies recorded for ≥15 days were classified as ‘treated’. Where no MRI information was recorded during a 3-month period, the value ‘unavailable’ was allowed. Where no new disability data were recorded during a 3-month period, the last previously recorded disability score was carried over.* At every time point, treatment status was a binary variable (treated/untreated) and each patient could contribute data to the treated and untreated pseudo-cohorts at different time points.

Relapses were recorded by treating neurologists, defined as new symptoms or exacerbation of existing symptoms persisting for ≥24 hours, in the absence of concurrent illness/fever, and occurring ≥30 days after a previous relapse. Disability was quantified with the Expanded Disability Status Scale (EDSS), excluding scores obtained <30 days after a relapse. Neurostatus EDSS certification was required at the participating centres.^20^

Presence/absence of new or enlarging T2 hyperintense lesions or contrast-enhancing lesions on cerebral MRI was reported by treating neurologists.

### Outcomes

The study endpoints were cumulative hazards of relapses, disability accumulation events and disability improvement events. Disability accumulation was defined as an increase in EDSS by 1 step (1.5 step if baseline EDSS=0 and 0.5 steps if baseline EDSS>5.5) confirmed by subsequent EDSS scores over ≥12 months. Disability improvement was defined as a decrease in EDSS by 1 step (1.5 steps if baseline EDSS≤1.5 and 0.5 steps if baseline EDSS>6) confirmed over ≥12 months, as over 80% of such events correspond to long-term accumulation of disability.^21^ No carryover EDSS scores were utilised in calculating confirmed disability endpoints. In addition, progression to EDSS step 6 confirmed over ≥12 months was evaluated in the primary analysis.

### Statistical analysis

Statistical analyses were conducted using R (version 3.0.3). In order to mitigate the effect of intermediates/confounders of treatment allocation and disease outcomes, marginal structural proportional hazards models were utilised.^22^ These models allowed comparison of counterfactual cumulative hazards of relapses, disability accumulation and disability improvement events between pseudo-cohorts never treated vs. treated with immunotherapies for 15 years from their first recorded visit.^23^

Marginal structural models estimated the probability of multiple events (Andersen-Gill^24^) with the partial likelihood function modified with inverse probability-of-treatment weights.^25^ Individual patient follow-up was right-censored at the last recorded EDSS score.

The stabilised non-normalised inverse probability-of-treatment weights were calculated at each 3-month period, using the ratio of the probabilities of treatment assignation conditional on baseline, time-dependent and stabilising variables (estimated with multivariable logistic regression models):^26^

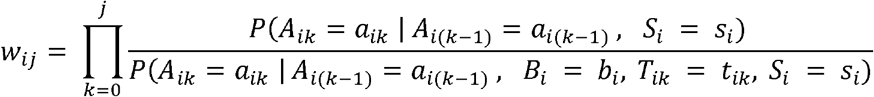

Here, *w_ij_* represents a stabilised weight for patient *i* at time *j. A* is the treatment status (treated/untreated) at time *k, S* represents stabilising variables (date of birth, date of first MS symptom, disease duration at first visit and pregnancy status at first visit), *B* represents baseline confounding variables and *T* represents time-dependent confounding variables (for the confounding variables see “Procedures”).

The weights reflect the probability of patients’ treatment status at any time depending on their demography and disease history, and were used to weigh contribution of a patient pseudo-cohort at any given period. Marginal structural models are inherently adjusted for all but the stabilising variables, including the history of the time-dependent variables.^25^ In addition to clustering by patient, the models were nested within study centre and right-censored at the last patients’ recorded EDSS. The primary analysis combined follow-up periods from the first to the last recorded disability score for each eligible patient, with time 0 defined as the beginning of the prospective follow-up (first clinic visit with a recorded EDSS). Each patient was allowed to contribute multiple treated and untreated periods to the analysis, depending on their treatment status at the given time (Supplementary Figure 1). Treatment was modelled as a time-dependent variable, relative to the time 0. For the summary of the study protocol see Supplementary Table 2. As a confirmatory analysis, we have repeated the primary analysis in a subset of patients followed for ≥15 years from their first recorded visit.

Tests of statistical inference were carried out at α=0.05. To assess stability of the associations estimated by the marginal structural models, non-parametric bootstrap with 1000 replicates was carried out.

### Sensitivity analyses

Seven sensitivity analyses were completed. Two analyses evaluated the effect of immunotherapy among patients with relapsing-remitting and progressive (primary and secondary) disease forms separately. A sensitivity analysis that examined the effect of segmentation of recorded follow-up into study periods was carried out by extending the study periods to 6 months. A sensitivity analysis among patients with complete follow-up from MS onset was carried out by restricting inclusion to the patients with their first recorded EDSS within the initial 3 months from the date of the first MS symptom. Another sensitivity analysis utilised the rigorously acquired prospectively recorded cohort from the MSBASIS sub-study, which requires prospective enrolment within 12 months from the first MS symptom and complete capture of EDSS functional system scores and MRI data.^27^ Finally, we have generalised the analysis to cohorts defined by disease duration and patient age by using two alternative definitions of baseline (time 0): the first recorded MS symptom (i.e. clinical onset of MS) and date of birth. This experimental approach was aimed at exploring the possibility of reconstructing disease trajectories over time expressed as MS duration or patient age among patients with incomplete follow-up and with leftside censoring.

### Data availability statement

The data analysed in this study are the property of the individual contributing centres. They can be made available upon reasonable request for the purpose of replication of the analyses included in this study and at the discretion of the principal investigators.

## Results

### Study population

Of the 34,007 patients included in the MSBase cohort as of 16/06/2015, 14,717 patients fulfilled the inclusion criteria and 1085 had ≥15 years of follow-up recorded from their first visit (Figure 1, Supplementary Table 4). The most common reason for exclusion was the lack of sufficient follow-up required for the analysis. A large proportion of these patients had only been enrolled in MSBase within 2 years prior to the database lock and have not yet accumulated sufficient data. The excluded patients tended to be captured later in their disease and with shorter prospective follow-up than the included patients (Table 2). Demographic information at first study visit was in keeping with the known epidemiology of MS (71% female, mean age 36 years, median disability EDSS step 2; Table 1). Median visit interval was 6 months, similar to most randomised clinical trials. Patients were exposed to immunotherapies for 69% of the prospectively recorded cumulative follow-up of 102,978 patient-years (median per-patient follow-up of 6 years). The most represented therapies were interferon β / glatiramer acetate (59% of follow-up time), followed by natalizumab (5%) and fingolimod (4%). The patients with ≥15-year follow-up were exposed to immunotherapies for 63% of the time over the median follow-up of 17 years. The time on higher-efficacy therapies was relatively less represented in this cohort compared to the full cohort, as natalizumab and fingolimod have only become available in 2006 and 2011, respectively.

**Figure 1.**
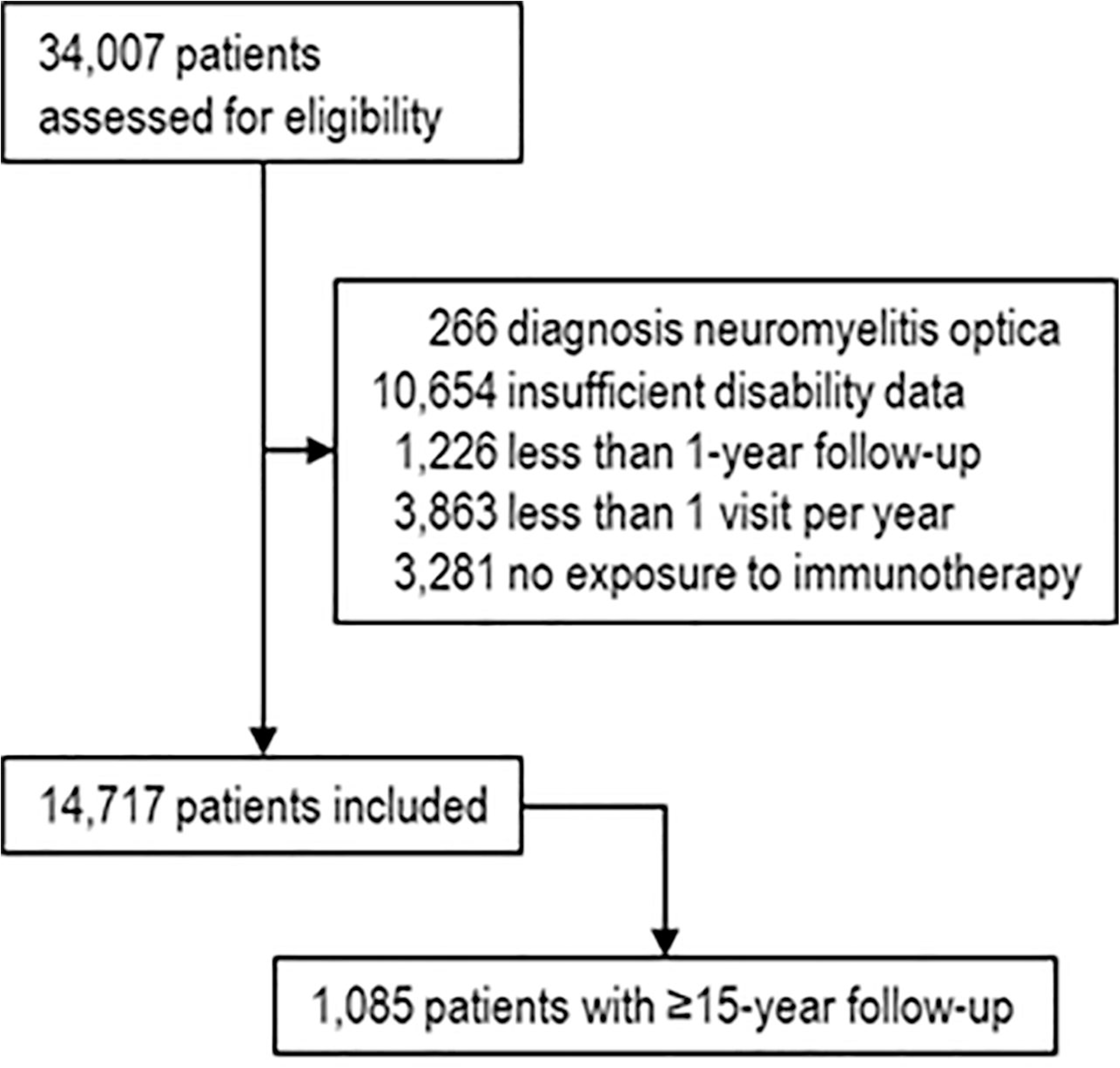
Patient disposition. The data quality procedure excluded 147 patient records: 95 from centres with less than 10 enrolled patients, 49 with missing birth date or the date of MS onset and 3 with erroneous information about disease progression. The inclusion criteria were applied so that patients’ follow-up is of sufficient duration to enable evaluation of at least short-term disability outcomes (≥1 year), with a minimum number of data points to ensure that individual hazard of confirmed disability worsening is non-zero (≥3 EDSS scores), sufficient data density to minimise the risk of disability events that were not captured and to minimise recall bias (≥1 EDSS score per year), The minimum data set was required for calculation of the inverse probability of treatment weights and outcomes. Patients had to be exposed to an MS immunotherapy at least once in order to eliminate indication bias, which is significant for untreated patients in countries where immunotherapies are commonly available. A large proportion of patients excluded from the analysis were enrolled in MSBase only within the prior 2 years and did not yet accumulate sufficient follow-up information.

**Table 1.**
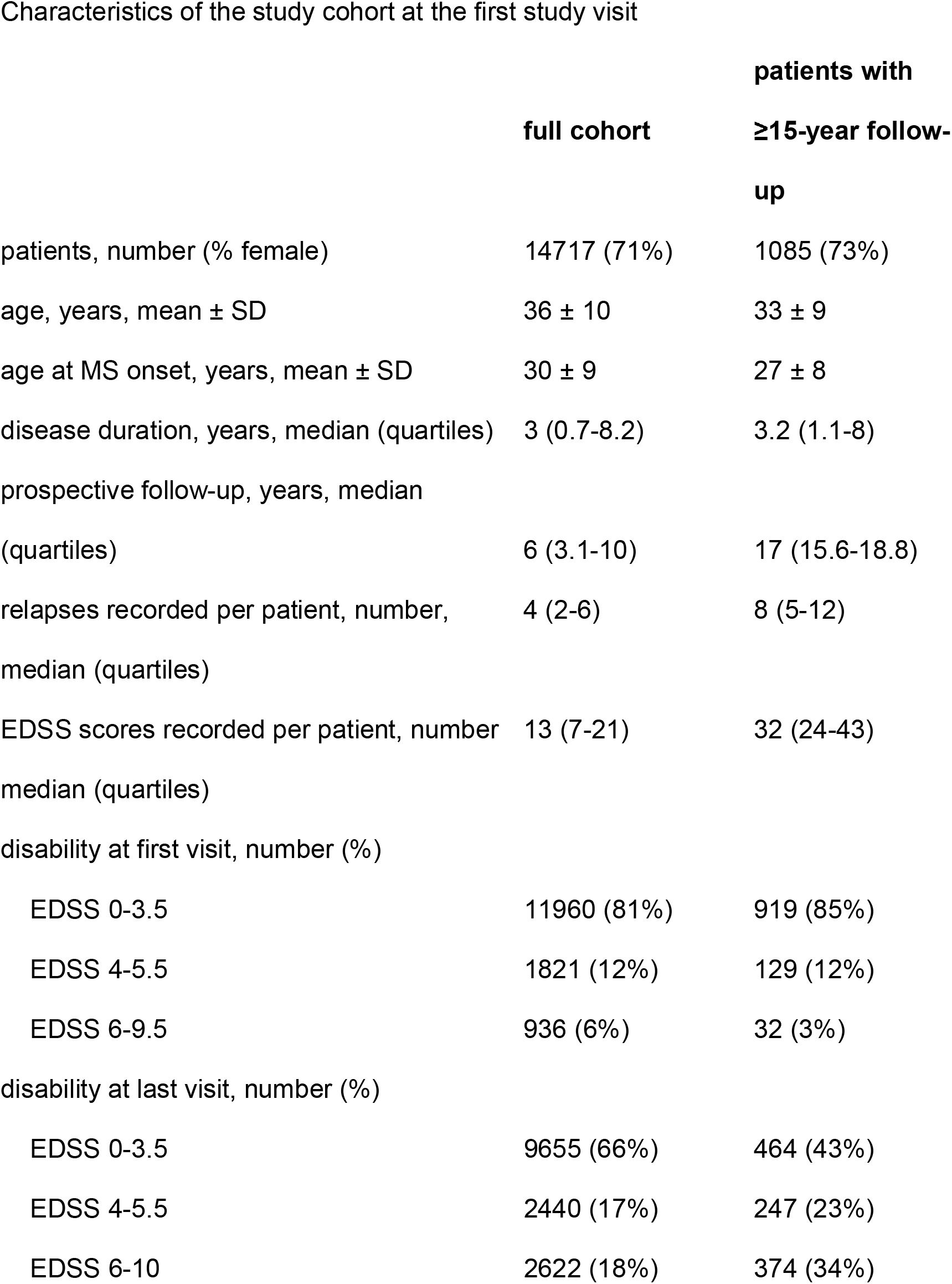
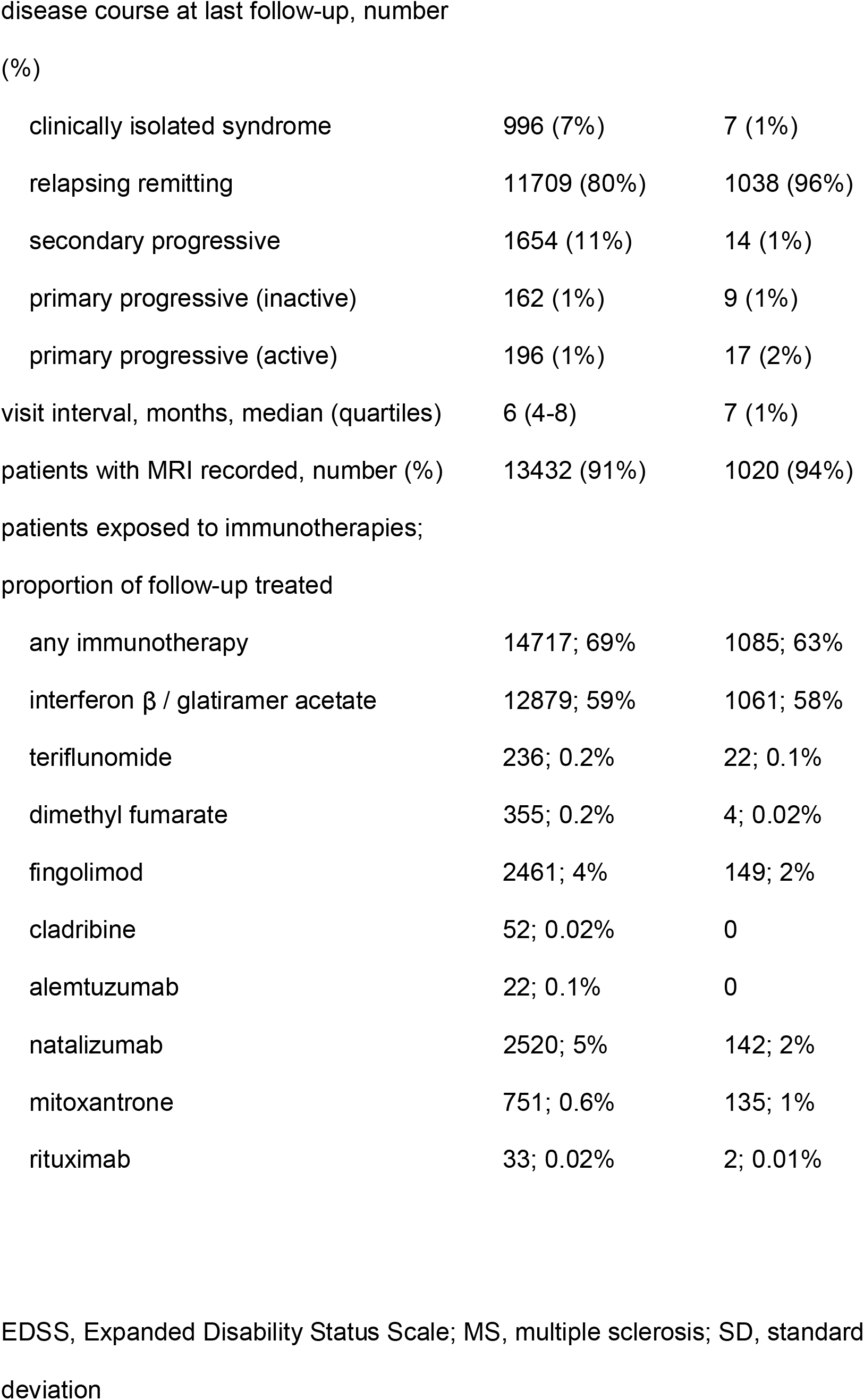
Characteristics of the study cohort at the first study visit

|  | **full cohort** | **patients with**<br>**≥15-year follow-**<br>**up** |
| --- | --- | --- |
| patients, number (% female) | 14717 (71%) | 1085 (73%) |
| age, years, mean ± SD | 36 ± 10 | 33 ± 9 |
| age at MS onset, years, mean ± SD | 30 ± 9 | 27 ± 8 |
| disease duration, years, median (quartiles) | 3 (0.7-8.2) | 3.2 (1.1-8) |
| prospective follow-up, years, median<br>(quartiles) | 6 (3.1-10) | 17 (15.6-18.8) |
| relapses recorded per patient, number,<br>median (quartiles) | 4 (2-6) | 8 (5-12) |
| EDSS scores recorded per patient, number<br>median (quartiles) | 13 (7-21) | 32 (24-43) |
| disability at first visit, number (%) |  |  |
| EDSS 0-3.5 | 11960 (81%) | 919 (85%) |
| EDSS 4-5.5 | 1821 (12%) | 129 (12%) |
| EDSS 6-9.5 | 936 (6%) | 32 (3%) |
| disability at last visit, number (%) |  |  |
| EDSS 0-3.5 | 9655 (66%) | 464 (43%) |
| EDSS 4-5.5 | 2440 (17%) | 247 (23%) |
| EDSS 6-10 | 2622 (18%) | 374 (34%) |
| clinically isolated syndrome | 996 (7%) | 7 (1%) |
| relapsing remitting | 11709 (80%) | 1038 (96%) |
| secondary progressive | 1654 (11%) | 14 (1%) |
| primary progressive (inactive) | 162 (1%) | 9 (1%) |
| primary progressive (active) | 196 (1%) | 17 (2%) |
| visit interval, months, median (quartiles) | 6 (4-8) | 7 (1%) |
| patients with MRI recorded, number (%) | 13432 (91%) | 1020 (94%) |
| patients exposed to immunotherapies; |  |  |
| proportion of follow-up treated |  |  |
| any immunotherapy | 14717; 69% | 1085; 63% |
| interferon $\beta$ / glatiramer acetate | 12879; 59% | 1061; 58% |
| teriflunomide | 236; 0.2% | 22; 0.1% |
| dimethyl fumarate | 355; 0.2% | 4; 0.02% |
| fingolimod | 2461; 4% | 149; 2% |
| cladribine | 52; 0.02% | 0 |
| alemtuzumab | 22; 0.1% | 0 |
| natalizumab | 2520; 5% | 142; 2% |
| mitoxantrone | 751; 0.6% | 135; 1% |
| rituximab | 33; 0.02% | 2; 0.01% |

EDSS, Expanded Disability Status Scale; MS, multiple sclerosis; SD, standard deviation

**Table 2.**
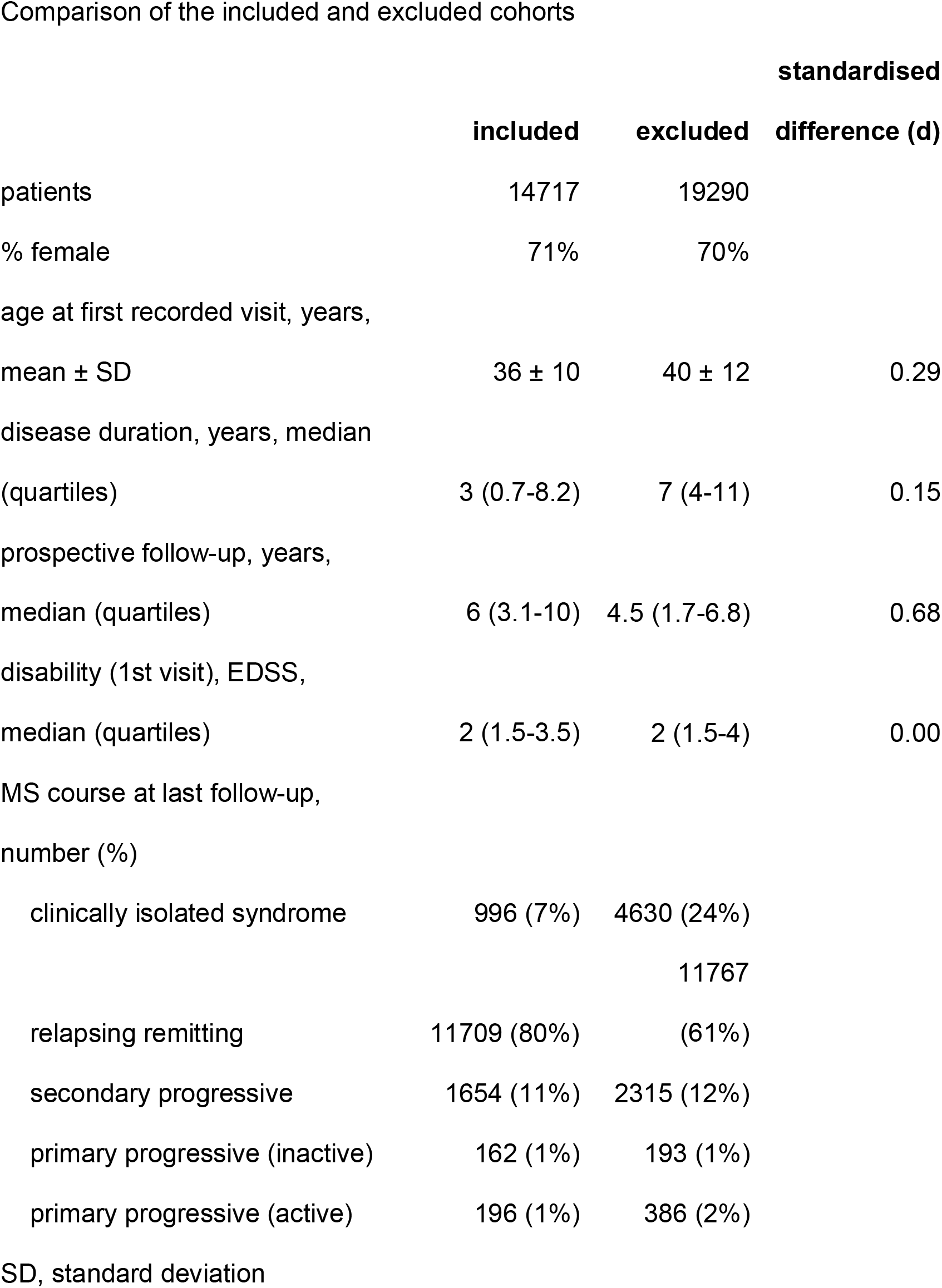
Comparison of the included and excluded cohorts

### Inverse probability-of-treatment weights

Stabilised non-normalised inverse probability-of-treatment weights for each patient and at each time point were built based on the probability of receiving immunotherapy at any given 3-month period conditional on patients’ demographic information, MS history and previous treatment exposure (for full list of baseline, time-dependent and stabilising variables, see Methods). The weights followed an expected distribution, centred around 1 and with only minor fluctuations over 15 years in both pseudo-cohorts, indicating good model specification (Supplementary Figure 3).

### Disease outcomes among all eligible patients

In the full study cohort, the pseudo-cohort treated continuously was less likely to experience relapses than the untreated pseudo-cohort (annualised relapse rate 0.32 vs. 0.46, respectively; hazard ratio [HR] 0.60, 95% confidence interval [95%CI] 0.43–.82, p=0.0016). Cumulative hazard of relapses in the treated vs. untreated cohorts was estimated at approximately 5 vs. 8 relapses at 15 years from first recorded visit, respectively (Figure 2). The difference between the treated and the untreated patients increased proportionally over time.

**Figure 2.**
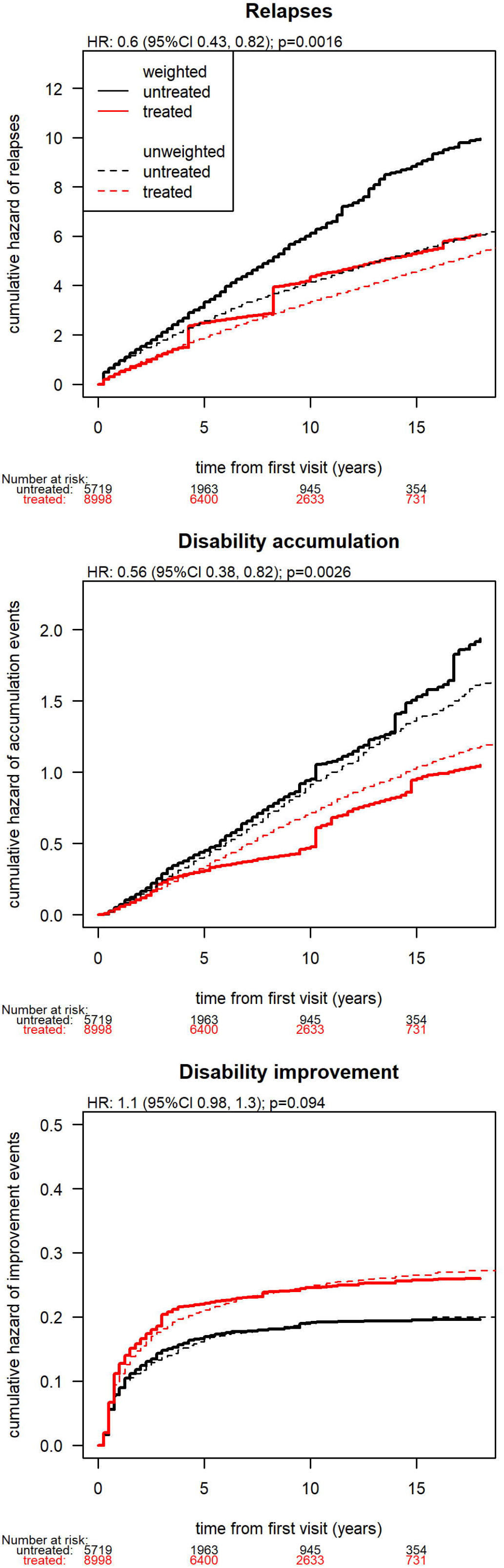
Incidence of relapses, disability accumulation and improvement in the treated and untreated pseudo-cohorts. Cumulative hazards for unadjusted models (dashed) and marginal structural models adjusted with inverse probability of treatment weights (solid) are shown. Numbers of patients contributing to the treated and untreated pseudo-cohorts are shown at multiple time points.

The treated cohort was less likely to experience 12-month confirmed disability accumulation events relative to the untreated cohort (HR 0.56, 95%CI 0.38-0.82, p=0.0026). Cumulative hazard of disability accumulation in the treated vs. untreated cohorts reached 1.0 vs. 1.5 events at 15 years, respectively (Figure 2). The difference between the treated and the untreated patients only became apparent at 3 years from first recorded visit and tended to increase with time. The bootstrap of the cumulative hazard of disability accumulation showed that the results of the primary analysis were robust to sampling variability (mean bootstrapped HR 0.57, standard error 0.06, Supplementary Figure 4).

The probability of reaching 12-month confirmed EDSS step 6 was markedly lower in the treated than the untreated cohort (HR 0.33, 95% CI 0.19-0.59, p=0.00019, Figure 3). Within 15 years from first visit, 13% of the treated cohort and 35% of the untreated cohort reached EDSS step 6.

**Figure 3.**
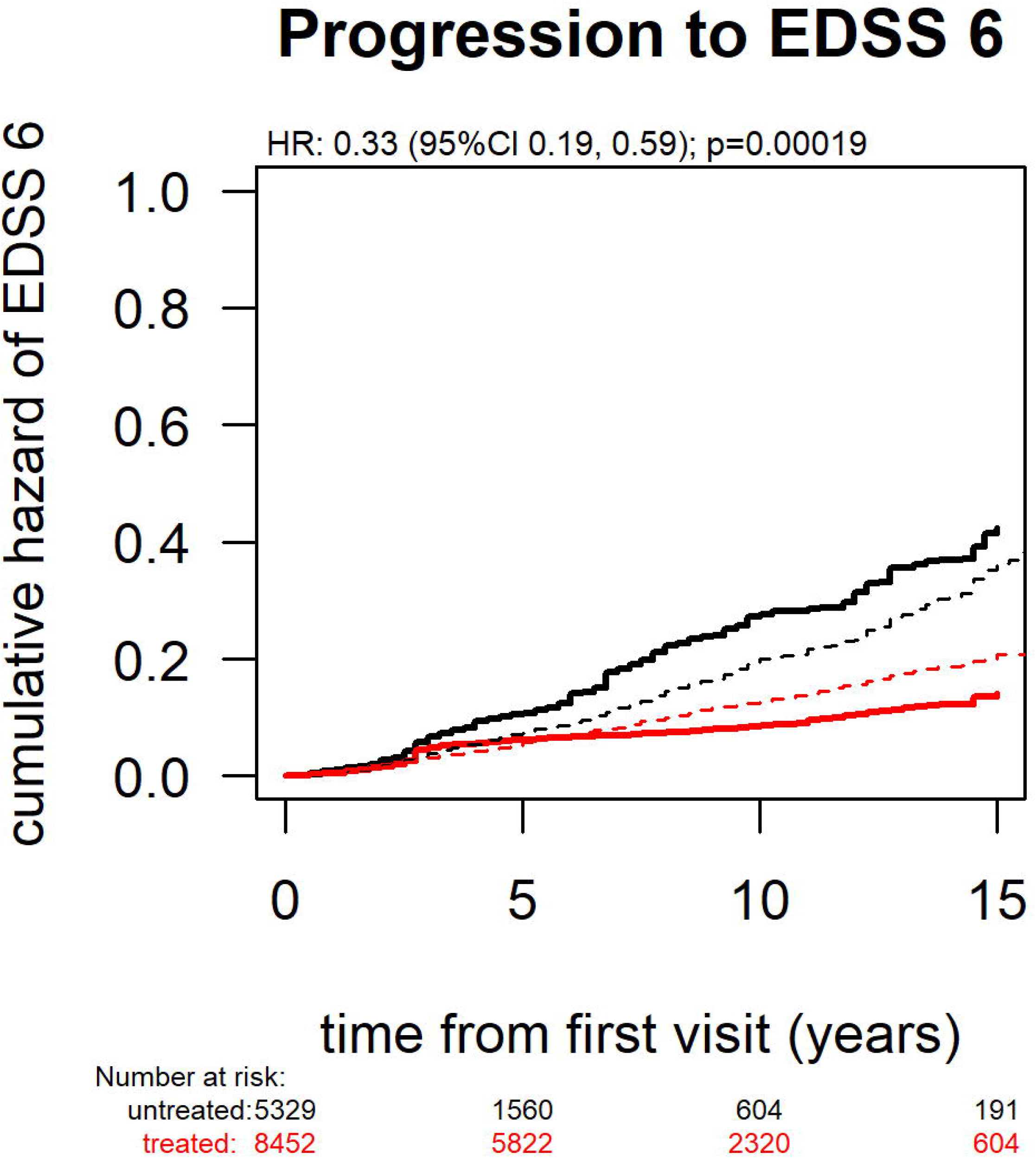
The risk of reaching EDSS 6 (patients use a single-point walking aid to walk ≥100 meters) in the treated and untreated pseudo-cohorts. Cumulative hazards for unadjusted models (dashed) and marginal structural models adjusted with inverse probability of treatment weights (solid) are shown. Numbers of patients contributing to the treated and untreated pseudo-cohorts are shown at multiple time points.

The probability of 12-month confirmed disability improvement events tended to be greater in the treated (0.21) than the untreated cohort (0.18) during the initial 4 years of follow-up. After year 4, the increment in the cumulative hazards was similar in the two pseudo-cohorts (Figure 2).Therefore, this trend did not reach the defined threshold for statistically significant difference (HR 1.10, 95%CI 0.98-1.30, p=0.094).

The analysis in relapsing-remitting MS-only confirmed the results of the primary analysis, demonstrating differences in relapse frequency (annualised relapse rate 0.35 vs. 0.56; HR 0.49, 95%CI 0.42-0.58, p<10^-16^), disability accumulation (HR 0.68, 95%CI 0.52-0.88, p=0.004) and disability improvement (HR 1.13, 95%CI 0.95-1.34, p=0.17). On the contrary, the analysis in progressive disease forms did not find any differences in disability accumulation (HR 0.92, 95%CI 0.81-1.05, p=0.22) or improvement (HR 1.38, 95%CI 0.85-2.26, p=0.19) between the treated and untreated pseudo-cohorts.

The sensitivity analysis using 6-month instead of 3-month study periods confirmed the results of the primary analysis. The treated cohort experienced lower frequency of relapses (annualised relapse rate 0.31 vs. 0.40, respectively; HR 0.54, 95%CI 0.49-0.59, p<10^-16^), lower hazard of disability accumulation (HR 0.74, 95%CI 0.62–0.88, p=0.0005) and greater probability of disability improvement than the untreated cohort (HR 1.26, 95%CI 1.11-1.42, p=0.0002).

### Disease outcomes among patients with ≥15-year follow-up

The results of the analyses among the 1085 patients with ≥15-year prospective follow-up from their first visit were in keeping with the results reported in the full study cohort. The treated cohort was less likely to experience relapses than the untreated cohort (annualised relapse rate 0.33 vs. 0.44, respectively; HR 0.59, 95%CI 0.50–0.70, p=10^-9^). Confirmed disability progression events were relatively less frequent in the treated cohort (HR 0.81, 95%CI 0.67-0.99, p=0.043). The probability of disability improvement did not differ between the treated and the untreated cohorts (HR 0.91, 95%CI 0.69-1.2, p=0.54).

### Disease outcomes among patients followed from disease onset

The sensitivity analysis that included only patients with first EDSS follow-up recorded ≤3 months after the first MS symptom identified 2194 eligible patients followed over 15,084 patient-years (69% female, mean age 31 years, median EDSS step 2, median follow-up 6 years, median visit interval 4 months). It replicated the results of the primary analysis for relapse incidence (showing lower hazard of relapses in the treated cohort, HR 0.51, 95%CI 0.38-0.68, p=10^-5^; Figure 4) and disability accumulation (lower hazard of disability accumulation in the treated cohort, HR 0.59, 95%CI 0.39-0.88, p=0.011). This sensitivity analysis also found a greater probability of disability improvement in the treated vs. untreated cohort (HR 1.36, 95%CI 1.02–1.80, p=0.038).

**Figure 4.**
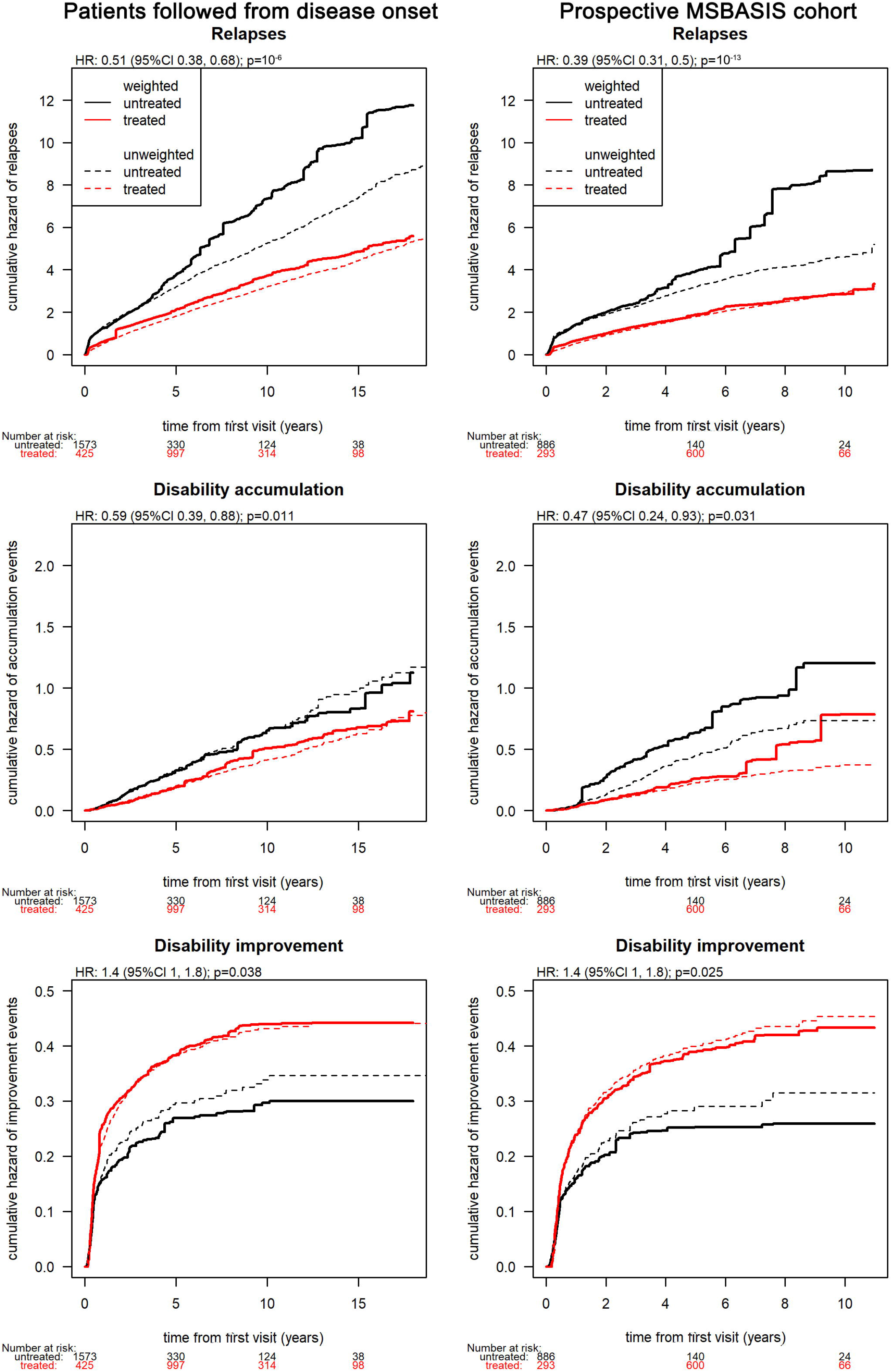
Sensitivity analyses: Comparisons of relapse frequency, disability accumulation and improvement between treated and untreated pseudo-cohorts consisting of patients followed from disease onset - i.e. with the first disability recorded within 3 months from first presentation of multiple sclerosis (left) and the prospective MSBASIS cohort (right). Numbers of patients contributing to the treated and untreated pseudo-cohorts are shown at multiple time points.

The sensitivity analysis utilising the prospective MSBASIS sub-study included 1291 patients followed over 7239 patient-years (69% female, mean age 31 years, median EDSS step 2, median follow-up 5.5 years, median visit interval 4 months). Similarly, this sensitivity analysis found superior outcomes in the treated cohort compared to the untreated cohort for relapse incidence (HR 0.39, 95%CI 0.31-0.50, p=10^-13^), disability accumulation (HR 0.47, 95%CI 0.24-0.93, p=0.031) and disability improvement (HR 1.36, 95%CI 1.04-1.79, p=0.025; Figure 4).

### Long-term disease outcomes throughout the duration of the MS and life span

When the time variable in the full study cohort was defined as the time from the first MS symptom (Supplementary Figure 5), the treated pseudo-cohort was less likely to experience relapses (annualised relapse rate 0.32 vs. 0.47; HR 0.54, 95%CI 0.4–50.65, p=10^-10^; Figure 5) and disability accumulation events (HR 0.69, 95%CI 0.55–0. 85, p=0.0007). No evidence of difference between the treated and the untreated pseudo-cohorts in disability improvement was found (HR 1.20, 95%CI 1.96-1.50, p=0.1).

**Figure 5.**
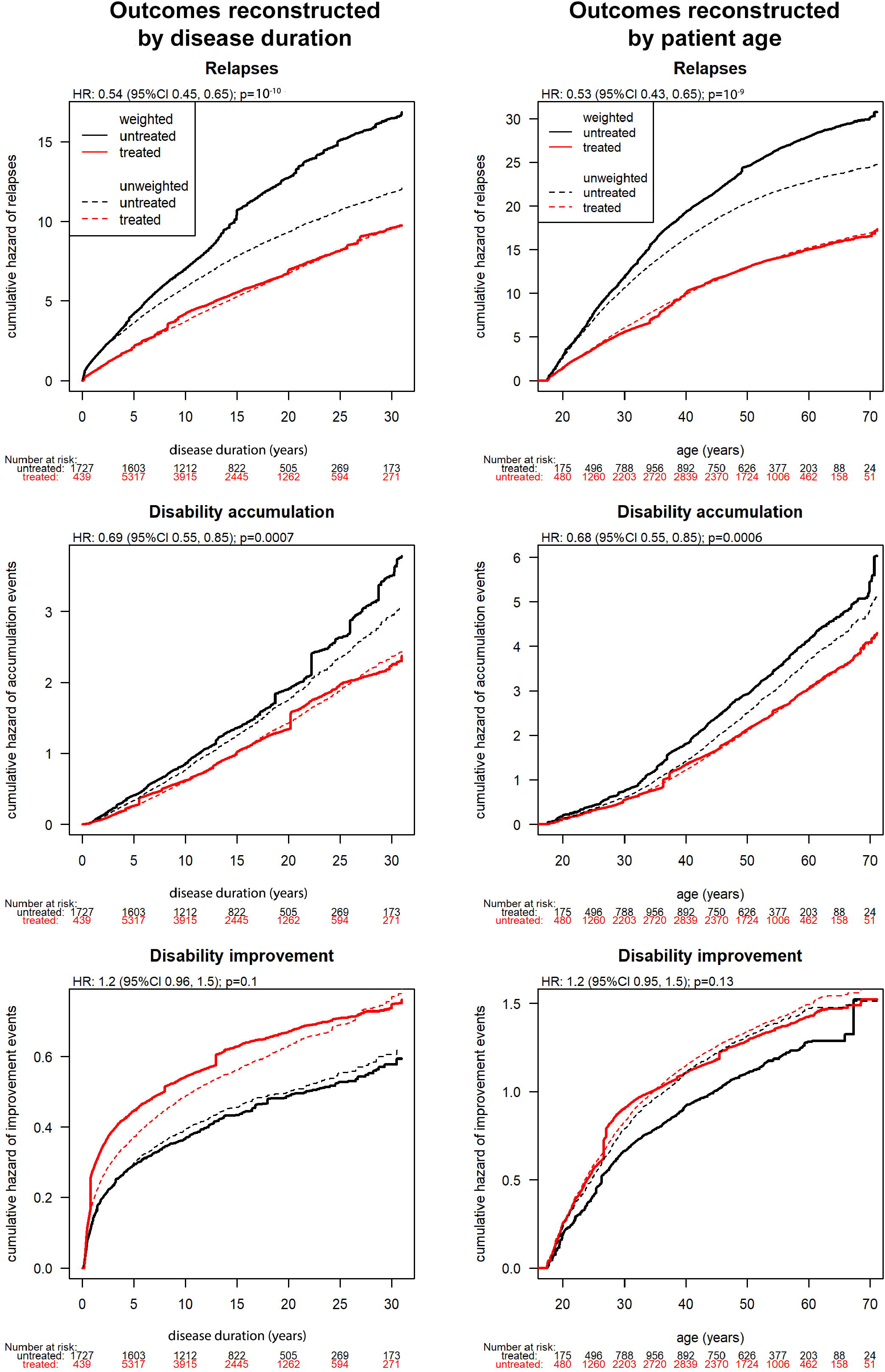
Sensitivity analyses: Generalisation of the analysis to follow-up by disease duration and patient age. Incidence of relapses, disability accumulation and improvement in the treated and untreated pseudo-cohorts analysed by disease duration and patient age. Cumulative hazards for unadjusted models (dashed) and marginal structural models adjusted with inverse probability of treatment weights (solid) are shown. Numbers of patients contributing to the treated and untreated pseudo-cohorts are shown at multiple time points.

Similarly, when the study follow-up was organised by patient age (Figure 5), the treated pseudo-cohort experienced a lower frequency of relapses (annualised relapse rate 0.32 vs. 0.46; HR 0.53, 95%CI 0.43-0.65, p=10^-9^) and disability accumulation events (HR 0.68, 95%CI 0.55-0.85, p=0.0006). The probability of disability improvement did not differ between the compared pseudo-cohorts (HR 1. 20, 95%CI 0.95-1.50, p=0.13).

## Discussion

### Principal findings

This observational study in 14,717 patients from the global MSBase cohort, including 1085 patients with ≥15-year recorded follow-up, demonstrated that continued immunotherapy reduces the risk of disability accrual in relapsing-remitting MS by 19–44% and the risk of impaired gait requiring use of a walking aid by 67% over 15 years.

The reduction in accumulation of disability was observed on the background of a 40–41% reduction in the frequency of MS relapses, an observation that is in keeping with the previous knowledge.^1^ Interestingly, we only observed a trend towards more likely improvement in disability in patients treated during early stages of MS. After a phase of accelerated disability improvement observed during the initial 4 years of follow-up, the probability of disability improvement became similar in the treated and untreated cohorts. This observation was confirmed statistically as a 36% higher chance of disability improvement among patients who were followed prospectively from disease onset.

### Comparison with other studies

In agreement with our conclusion, several previous studies showed reduced disability during treatment with immunotherapies.^2, 5^ The results of our study are supported by a study of 5,610 patients enrolled in the UK Risk Sharing Scheme, which showed, using Markov and multilevel continuous models, that treatment with interferon β or glatiramer acetate was associated with a 24% decrease in disability accrual over up to 6 years.^13, 14^ An MSBase study among 2,466 patients with relapsing-remitting MS suggested that continuous exposure to interferon β or glatiramer acetate for 10 years was associated with a mean mitigation of disability accrual by 0.86 EDSS steps.^28^ Re-assessment of the pivotal trial of interferon β-1b at 16 years (n=372), using recursive partitioning, showed that earlier exposure to interferon β was associated with a decreased hazard of reaching disability milestones or death.^9^ A propensity score-weighted analysis in 1,504 patients showed that patients treated with interferon β were less likely to reach EDSS step ≥4 (restricted ambulatory capacity) than untreated patients over up to 7 years.^29^ In keeping with our previous studies, we did not observe an overall effect of pooled immunotherapies in non-selectively treated cohorts with progressive disease forms.^30^ However, this observation does not rule out the possibility that some therapies can slow down progression of disability in progressive disease phenotypes with superimposed episodic inflammatory activity.^31^

In contrast to our results, a propensity score-matched analysis in 2,656 patients did not find any evidence of difference in the probability of reaching EDSS step ≥6 (use of unilateral support to walk ≥100 meters) between patients treated with interferon β and those untreated. Interestingly, a trend favouring the treated cohort was observed when compared with historical controls, while an opposite trend was seen in the comparison against contemporary controls.^12^ In the ensuing debate, the conflicting trends were attributed to the residual selection bias and the lack of re-adjustment for the ongoing decision process of choosing between treatment and no treatment, which occurs continuously throughout long-term follow-up. This problem of continuous confounding of treatment allocation was also inherent in the other prior comparisons of longitudinal outcomes between treated and untreated patients.

### Methodological considerations

The key problem in uncovering unbiased causal effect of immunotherapy on disability accrual is therefore that of time-dependent confounding.^23^ Karim and colleagues used a marginal structural Cox model^32^ applied to a clinic-based cohort from British Columbia to assess the association between treatment and disability accrual. Using marginal structural models allowed the authors to construct pseudo-cohorts defined by their treatment status at any time point and re-adjusted for time-dependent confounders and intermediates of treatment allocation and study outcome.^33^ The study demonstrated how marginal structural models can effectively mitigate time-dependent confounding, and did not find an association between treatment with interferon β and the risk of reaching EDSS step ≥6. In our present study, we have extended the methodology used by Karim et al. to enable inclusive analysis of all treated patients, whereby the clinical follow-up in every patient consists of both treated and not treated periods. In the presented models, every patient may contribute such periods to either treated or untreated group at different times – defined with respect to their first recorded visit, disease onset or date of birth. Naturally, it is not possible to directly observe the outcomes of two mutually exclusive treatment decisions in a single cohort; therefore we have utilised a counterfactual framework which uses a well-defined statistical methodology, and have termed the compared groups ‘pseudo-cohorts’.^13, 15, 22^ This approach enabled us to compare cumulative hazards of disability and relapse events in pseudo-cohorts that were hypothetically treated or untreated for 15 years from their first visit (or throughout their disease duration or life span). The assumption of such approach is that the effect of therapy does not attenuate over time. We have observed that the effect of therapy on relapses and worsening of disability was sustained throughout the 15-year follow-up (see Figures 2 and 3). This observation supports continued treatment with immunotherapies over an extended period of time. In contrast, the trend towards a difference in disability improvement was only restricted to the initial years following the first presentation of MS. This phenomenon is likely associated with functional compensatory mechanisms, which become exhausted with increasing cumulative inflammatory damage and age.^34^ Such observation supports the notion that in order to facilitate recovery of neurological function immunotherapies should be commenced during early stages of MS, when a recovery from disability is relatively more likely.

To fulfil the concept of a study cohort as a ‘group that shares defining characteristics’, the outcomes in all patients were analysed from their first recorded visit. Thus, the primary analysis did not require left-censoring. In order to ensure stability of the observed associations over the long-term, we have replicated the primary analysis in a sub-cohort with ≥15 years of recorded continuous follow-up. Furthermore, we have explored presentation of the hazards of disease outcomes in the context of disease duration or patient age. Using the first clinical presentation of MS or birth date as study baseline, respectively, these two sensitivity analyses have replicated the results of the primary analyses in full. While this suggests that combining observed periods to reconstruct disease outcomes over time that was not observed continuously is a feasible strategy, further mathematical justification of such approach is required. In order to eliminate the scenario when treated and untreated cohorts are not comparable (due to strong indication bias that would lead to the allocation of patients with ‘benign MS’ to the untreated cohort) and to fulfil the assumption of positivity, we have restricted inclusion to only those patients who qualified for at least one immunotherapy during the course of their disease.^35^ The study cohort was exposed to injectable therapies for 59% of the studied time, to more potent therapies for 9% of the time and was untreated for 31% of the time. Within the treated cohort, sequencing of therapies driven by clinical reasoning could lead to maximising the true treatment effectiveness when compared with efficacy reported from randomised blinded trials, as expected.^36^ The marginal structural models enabled us to draw inference about the causal associations between current treatment status and the probability of recurrent disability and relapse events, while accounting for measured confounders and intermediates of treatment allocation and disease outcomes.^37^ Thus, the results presented can be considered free from measurable bias of treatment assignation.

### Limitations

The main limitation of this study is inherent in the observational nature of the analysed data that originated from a large multicentre clinical cohort. We have mitigated the impact of inter-centre variability by applying a rigorous data quality procedure and nesting the models within study centres. The generalisability of the presented results may be restricted to patients followed in academic MS centres. While we have extended the analytical methodology to enable comparison of cumulative hazards of recurrent events, in its present form, the models do not allow us to evaluate delayed effects (i.e. delayed disability worsening and improvement or advancement to secondary progressive disease stage, whose risk could be modulated by treatment exposure in the immediate or distant past) or directly compare outcomes between multiple therapies. However, the models were adjusted for prior treatment status (whether immediately prior to each 3-month period or the overall cumulative treatment history), relapses, disability accumulation and improvement. Due to our inclusive definition of treated period (≥15 days on treatment), classification of immunosuppressants as untreated periods, and pooling of low-and high-efficacy immunotherapies, the true differences between the treated and untreated pseudo-cohorts may be underestimated. It is therefore reassuring that clinically meaningful and consistent differences in relapse and disability outcomes were shown despite our conservative study design. While we have mitigated the risk of measured confounding by including a large number of potential confounders in the models of inverse probability-of-treatment weights and by completing seven sensitivity analyses, propensity score-based methods can be subject to unmeasured confounding. Finally, many of the assumptions of marginal structural models are unmeasurable. Therefore, consistency of the results across the primary and the sensitivity analyses - among patients with different definitions of baseline and study period - provides additional assurance with respect to the robustness of the used models.

### Conclusions and implications

This study provides class III evidence that long-term exposure to immunotherapy not only reduces relapse activity but also prevents at least a fifth of neurological disability worsening in patients with relapsing-remitting MS. In early MS, accelerated recovery from previously accrued disability can be observed early after commencing immunotherapy. This information is highly relevant to the therapeutic decision process, highlighting the long-term, clinically meaningful benefits of early and continued immunotherapy on preserving patients’ physical capacity.

## FOOTNOTE

*EDSS was recorded during two consecutive 3-month periods at 84,226 time points. These EDSS values were highly correlated, with r=0.95.

## ACKNOWLEDGEMENTS

We thank the patients and their carers who have agreed to participate in the global MSBase cohort study. The list of MSBase Study Group co-investigators and contributors is given in Supplementary Table 1.

This study was financially supported by National Health and Medical Research Council of Australia [1129189, 1140766, 1080518] and Biogen [research grant 2016003-MS]. The MSBase Foundation is a not-for-profit organization that receives support from Roche, Merck, Biogen, Novartis, Bayer-Schering, Sanofi-Genzyme and Teva. The study was conducted separately and apart from the guidance of the sponsors.

## Appendix 1: Author contributions

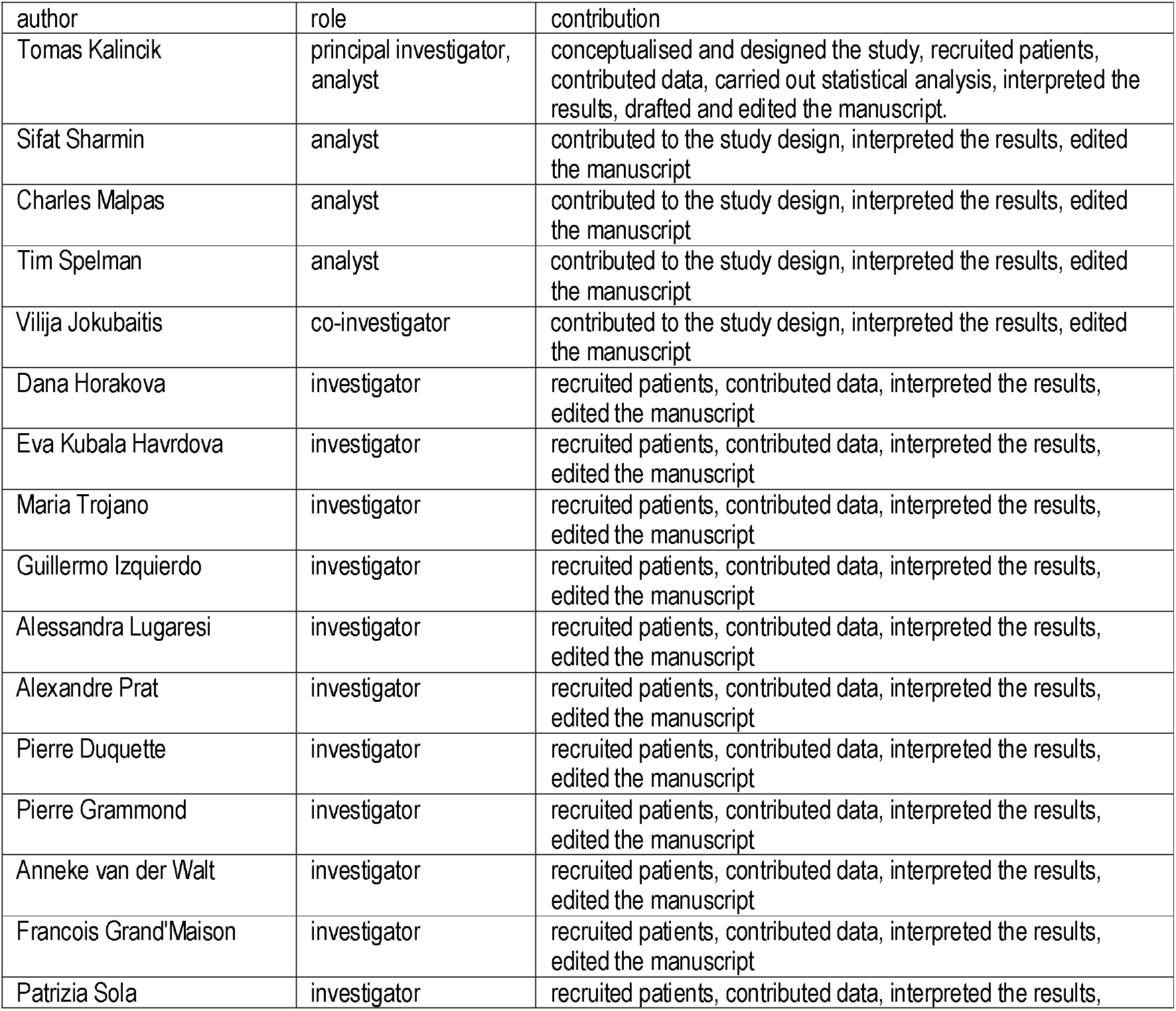
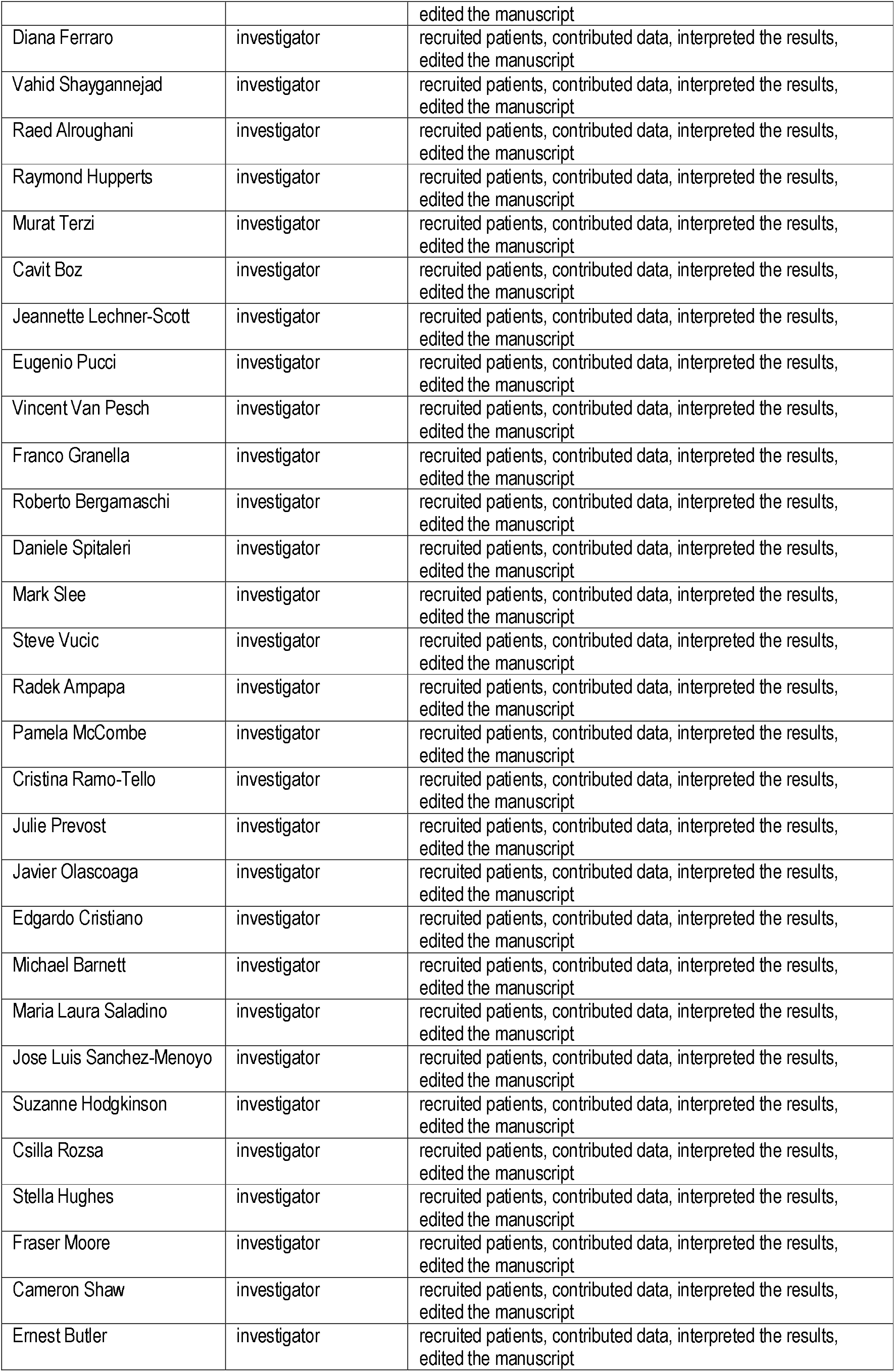
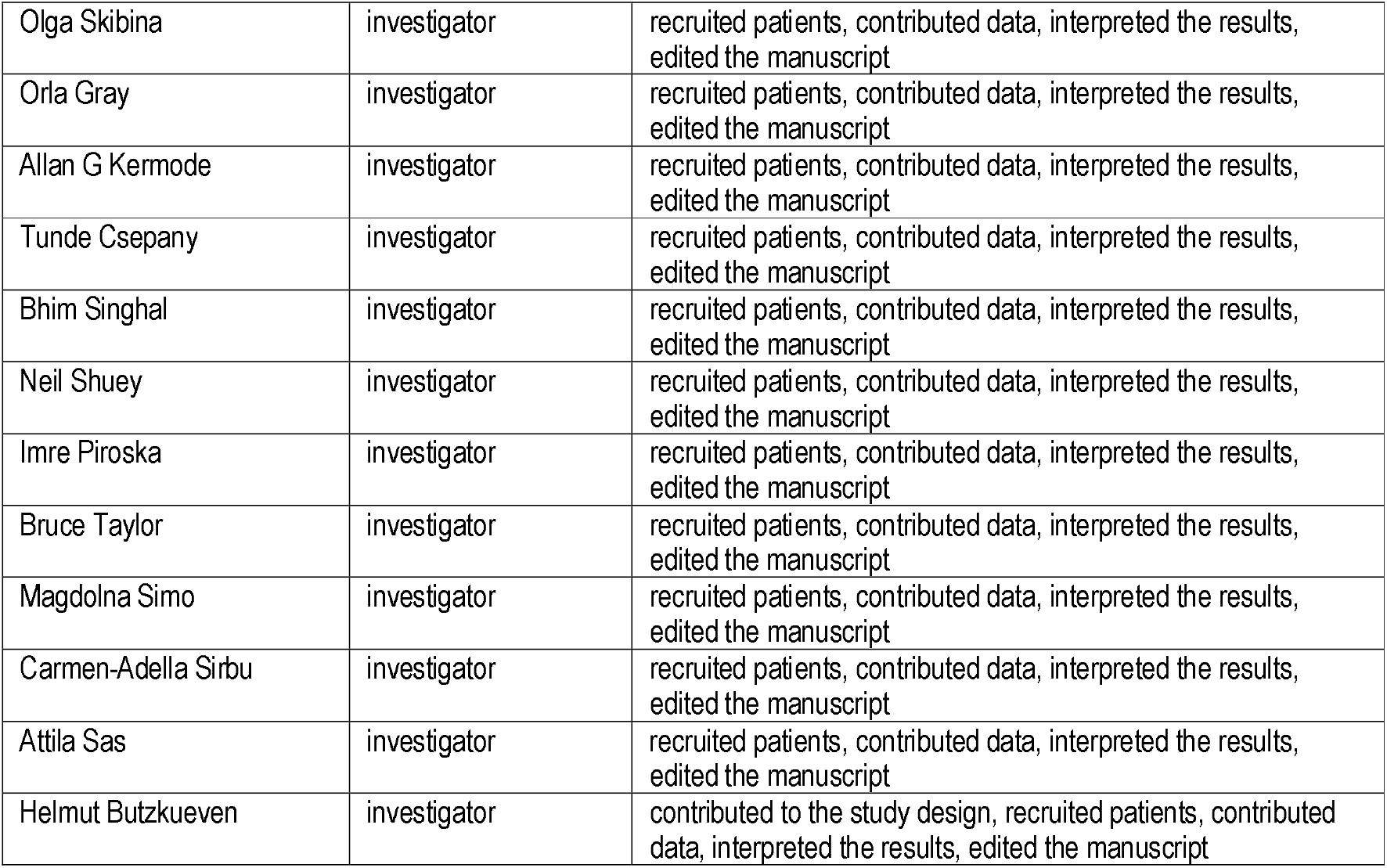

## Appendix 2: Co-investigators

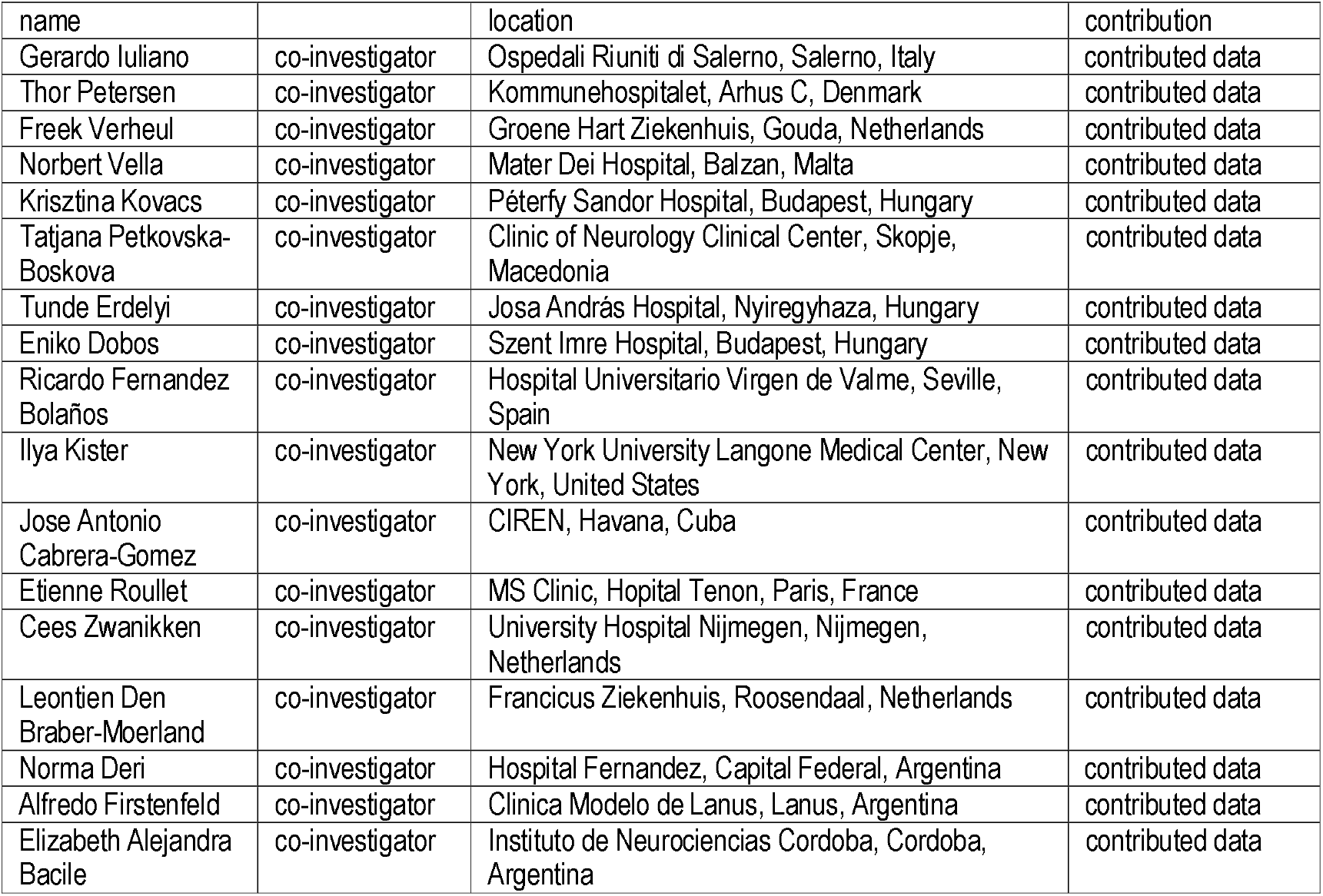
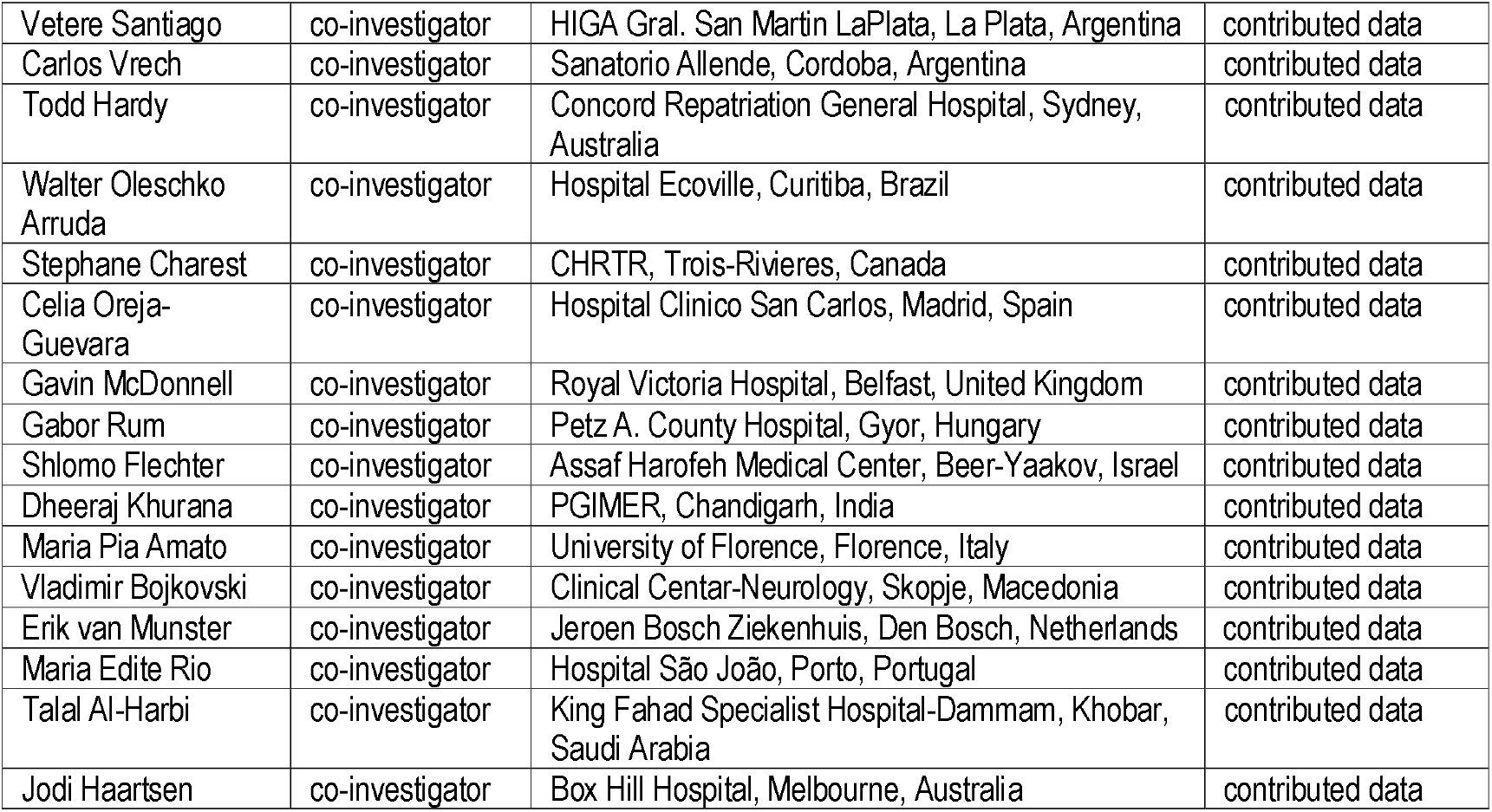

